# Rac1 Activated by DOCK1 in Combination with Rab31 Promotes the Development of Diabetic Retinopathy

**DOI:** 10.1101/413443

**Authors:** Guijun Xie, Peilin Lv, Man Li, Lingju Kong

## Abstract

**Background:** Diabetic retinopathy (DR) is one of the most common and severe microvascular complications of diabetes. It’s a fundus lesion with specific changes, and its specific molecular mechanism is still unclear.

**Methods:** All target proteins and markers expression in the study was verified by qPCR and western bloting. The morphology and behavior of human retinal epithelial cells ARPE-19 were analyzed using immunofluorescent and apoptosis detection assays, respectively. And, Small GTPase activity was detected by the GST-pull assay.

**Results:** We found that DOCK1 showed high expression in high glucose-induced DR. Moreover, its high expression is positively correlated with Rab31 in tissues. DOCK1 promotes the activation of Rac1 and promotes apoptosis of retinal epithelial cells. Rab31 promotes the degradation of TJ proteins by promoting the transport of TJ proteins from the plasma membrane to the endosomes, thereby affecting the tight junctions of epithelial cells. Inhibition of Rac1 activity can effectively reduce cell apoptosis. Similarly, inhibition of Rab31 activity can significantly inhibit the degradation of TJ proteins.

**Conclusion:** This study based on the high glucose-induced DR cell model reveals the role of two mutually synergistic signaling pathways through the important cytological phenomena of apoptosis and damage of tight junctions. One is the degradation of TJ proteins caused by Rab31-mediated vesicle trafficking, and the other is the apoptosis of retinal epithelial cells mediated by DOCK1-Rac1.

## 1. Introduction

Diabetic retinopathy (DR) is one of the most common and severe microvascular complications of diabetes [1]. Clinically, according to the presence or absence of retinal neovascularization, DR without retinal neovascularization is called non-proliferative diabetic retinopathy (NPDR) (or simple or background type), and DR with retinal neovascularization is called proliferative diabetic retinopathy (PDR) [2]. DR is a fundus lesion with specific changes, and its specific molecular mechanism is still unclear.

In the early stages of DR, retinal microvascular endothelial cell (RMVECs) barriers are first impaired, such as tight junctions (TJ), adhesion junctions (AJ), gap junctions (GJ), and complex adhesions (CA) [3]. In all of these biomarkers of endothelial dysfunction, TJ have a high abundance in RMVEC. Other evidence suggested that endothelial cell TJ damage may be an important mechanism for increased endothelial cell permeability [4]. However, the molecular mechanism of endothelial cell TJ damage in DR patients has not been clearly studied. Studies based on other disease models have revealed that some small GTPases may play an important role in the process of TJ endocytic transport, digestion and degradation, such as Rab family small GTPases which regulates intracellular vesicle trafficking and Rho family of small GTPases which regulates intracellular cytoskeleton dynamic changes in the [5,6].

As a small GTPase of the Rho family, Rac1 plays an important role in many intracellular signaling pathways [7-9]. In many cancers, activation of Rac1 activates downstream ERK1/2 and AKT signaling pathways, promotes the cytoskeleton dynamic changes in tumor cells, leading to proliferation or migration, and ultimately promotes tumor metastasis [10,11]. Studies have shown that the Tiam1-Rac1-Nox2 signal axis was activated during the initial stages of diabetes to increase intracellular reactive oxygen species (ROS), leading to mitochondrial damage and accelerated capillary cell apoptosis [12]. Activation of Tiam1-Rac1-mediated Nox2 and p38 MAP kinases constitutes an early signaling event leading to mitochondrial dysfunction and DR development [13]. Recently, it has been reported that in diabetes, acetylation of NF-κB mediates Rac1 transcriptional activation in the retina, and the regulation of acetylation in the early stages of DR has the potential to inhibit its development [14]. These indicate that Rac1 plays an important role in the development of DR from oxidative damage and transcriptional regulation to cell proliferation and apoptosis. However, as a small GTPase, its activation requires guanine nucleotide-exchange factor (GEF) regulation. In addition to the above Tiam1, DOCK1 is also a typical GEF for Rac1 activating, which also plays an important role in the occurrence and development of tumors [15]. However, there is no relevant research in DR.

Different from Rac1, there are few studies on the small GTPases of Rab family in the development of DR. Studies have shown that key signaling pathways involved in Wnt-MAPK signaling pathways or neuroinflammation are controlled by epigenetics in fibrotic disorders involving retinal detachment, and the results also emphasize neurovascular formation (ETS1, HES5, PRDM16) contribution in DR. The authors also pointed out the role of Rab31 in epigenetic regulation in this study [16]. In addition, the Rab family plays an important role in the regulation of TJ. Studies have shown that Rab13, Rab14 and Rab27 play a role in the regulation of tight junctional structures in cells [5-6,17].

Based on database analysis, this study found that DOCK1 showed significant high expression in PDR vascular membrane tissue, which also indicates that Rac1 may play an important role in the development of DR. At the same time, according to the analysis of the database, we also found that the small GTPase Rab31, which regulates intracellular vesicle trafficking in PDR vascular membrane tissue, also showed a high expression trend. There are few reports on the progress of two small GTPases in the coordinated regulation of DR. Here, their specific regulatory mechanisms in the cell model of DR were studied, revealing their specific molecular mechanisms and providing a theoretical basis and possible molecular targets for the treatment of DR.

## 2. Materials and methods

### 2.1 Cell line and Cell culture

The human retinal epithelial cells ARPE-19 cells were purchased from the ATCC Biological Resources Center. After resuscitation cells were cultured in Dulbecco’s modified Eagle’s medium (DMEM) containing 10% fetal bovine serum (FBS) (Gibco, Life Technologies), 100 U/ml penicillin, and 100mg/ml streptomycin. The cells were incubated in a humidified atmosphere of 95% air and 5% CO2 at 37°C.

### 2.2. Plasmids and siRNAs

Full-length DOCK1 and Rac1 wese amplified from cDNA. The polymerase chain reaction (PCR) products were cloned into the pCMV-N-Flag (Beyotime, Nantong, China). The cells were seeded in 6-well plates, cultured to 80~90% confluence, and then transiently transfected with the plasmid by using Lipofectamine 3000 (Invitrogen) according to the reverse transfection method provided by the manufacturer.

Duplex oligonucleotides were chemically synthesized and purified by GenePharma (Shanghai, China). The small interfering RNA (siRNA) duplexes used were DOCK1, #1, 5’-UUUUCCAUUCCUUUCGAGCCG-3’, Rac1, #1, 5’-AGGAAAACUGCAGAGAAACAG-3’ and Rab31, #1, 5’-UGAUGUUGUGGUCAAAGUGAU-3’. Cells were transfected with siRNA duplexes using Lipofectamine 3000 (Invitrogen) according to the reverse transfection method provided by the manufacturer.

### 2.3 Real time-qPCR

The collected cells were prepared for total RNA extraction using using Trizol reagent (Invitrogen, USA). The cDNAs were synthesized using a reverse transcription kit (Quant One Step RT-PCR kit, TIANGEN, China), and real-time PCR was performed using SYBR green master mix (Quant one step qRT-PCR Kit, TIANGEN, China) on a MyiQTM2 (BIORAD, USA). GAPDH was used to normalize the real-time PCR data. The following primer sequences were used: DOCK1, 5’-ACCGAGGTTACACGTTACGAA-3’ and 5’-TCGGAGTGTCGTGGTGACTT-3’ Rab31,5’-GTGCCTTCTCGGGGACAC-3’ and 5’-TGCCCCAATAGTAGGGCTGA-3’GAPDH, 5’-GGAGCGAGATCCCTCCAAAAT-3’ and 5’-GGCTGTTGTCATACTTCTCATGG-3’.

### 2.4 Western blotting

The cells were lysed in radioimmune precipitation assay (RIPA) buffer supplemented with protease inhibitor cocktail (Roche, Shanghai, China) to prepare the protein sample. Protein concentration was measured using Enhanced BCA Protein Assay Kit (Beyotime Biotechnology, China) and 40µg of protein were loaded and separated by 10% sodium dodecyl sulfate polyacrylamide gel electrophoresis (SDS-PAGE). Proteins were transferred to NC membranes and then blocked with 5% BSA in Tris-buffered saline with Tween-20 before detection with the following antibodies: Occludin (1:200, Santa Cruz, USA), ZO-1 (1:1000, CST, USA), Claudin-5 (1:200, Santa Cruz, USA), DOCK1 (1:1000, CST, USA), Rab31 (1:200, Santa Cruz, USA), Rac1 (1:1000, CST, USA), EEA1 (1:1000, CST, USA), M6PR (1:1000, CST, USA), LAMP1(1:1000, CST, USA), GAPDH (1:1000, CST, USA). After Primary antibody incubation for 12 hours, the NC membranes were washed by TBST buffer before incubated with secondary antibody for 1h. Specific binding was visualized by ECL reaction.

### 2.5 Immunofluorescent staining

Cells were fixed with 4% paraformaldehyde (PFA). Then, the cells were incubated with primary antibody against Occludin (1:50, Santa Cruz, USA), Rab31 (1:50, Santa Cruz, USA), EEA1 (1:200, CST, USA), M6PR (1:200, CST, USA) or LAMP1 (1:200, CST, USA), followed by washing and incubation with AF594-conjugated goat anti-rabbit secondary antibodies (1:250; Earthox, USA) for 2h at 37°C. The nuclei were stained with 4’, 6-diamidino-2-phenylindole (DAPI). Fluorescent images were visualized and captured using an inverted fluorescence microscope (Olympus, Tokyo, Japan). Images were visualized and captured using a phase contrast microscope (Olympus, Tokyo, Japan).

### 2.6 Apoptosis detection assay

Cells were seeded in a 96-well plate as a density of 5000 cells/well for 24 h. Cell apoptosis was measured using the Annexin V-FITC/PI Apoptosis Detection Kit (Ye Sen Biological Technology, China), according to the manufacturer’s instructions.

### 2.7 GST-pulldown assay

Glutathione S-transferase (GST)-tagged GTP-Rac1 binding protein (RBP) and GTP-Rab31 binding protein (RBP31) fusion proteins were expressed in Escherichia coli. Each purified GST fused proteins were immobilized on 40 μl of Glutathione-Sepharose 4B beads (GE Healthcare, USA) and equilibrated in the pull-down binding buffer consisting of 50 mM HEPES, 50 mM NaCl, 0.1% NP-40, and protease inhibitors. The purified complex was added to each affinity beads and the binding reactions were incubated at 4°C for 3 h. The beads were washed three times using the binding buffer. The bound proteins were analyzed by SDS-PAGE followed by Coomassie staining.

### 2.8 Statistical Analysis

Data are presented as means ± SEM of several experiments. Statistical comparisons were performed by either Student’s 2-tailed t test or ANOVA with Tukey multiple comparison post-test, as appropriate. A P value of less than 0.05 was considered significant.

## 3. Results

### 3.1 High expression and correlation of DOCK1 and Rab31 in DR

To determine the role of DOCK1 and Rab31 in DR progression, we analyzed the association of DOCK1 and Rab31 mRNA expression in two DR patient-related sample data sets (GSE53257 and GSE60436) and two streptozotocin (STZ)-induced diabetic mouse model sample data sets (GSE24423 and GSE20886). The results showed that mRNA expression levels of DOCK1 and Rab31 were increased in FVMs of DR patients compared with normal ocular fibrovascular membrane (FVMs) (DOCK1: *P*<0.0001; Rab31: *P*=0.006) (Figure 1A and B). Furthermore, we evaluated the association between DOCK1 and Rab31 expression in tissues. There was a positive correlation between DOCK1 and Rab31 mRNA expression in DR-related samples (r=0.521, *P*=0.002) (Figure 1C). These results all suggested that DOCK1 and Rab31 may play a important role in the development of DR.

**FIG.1.**
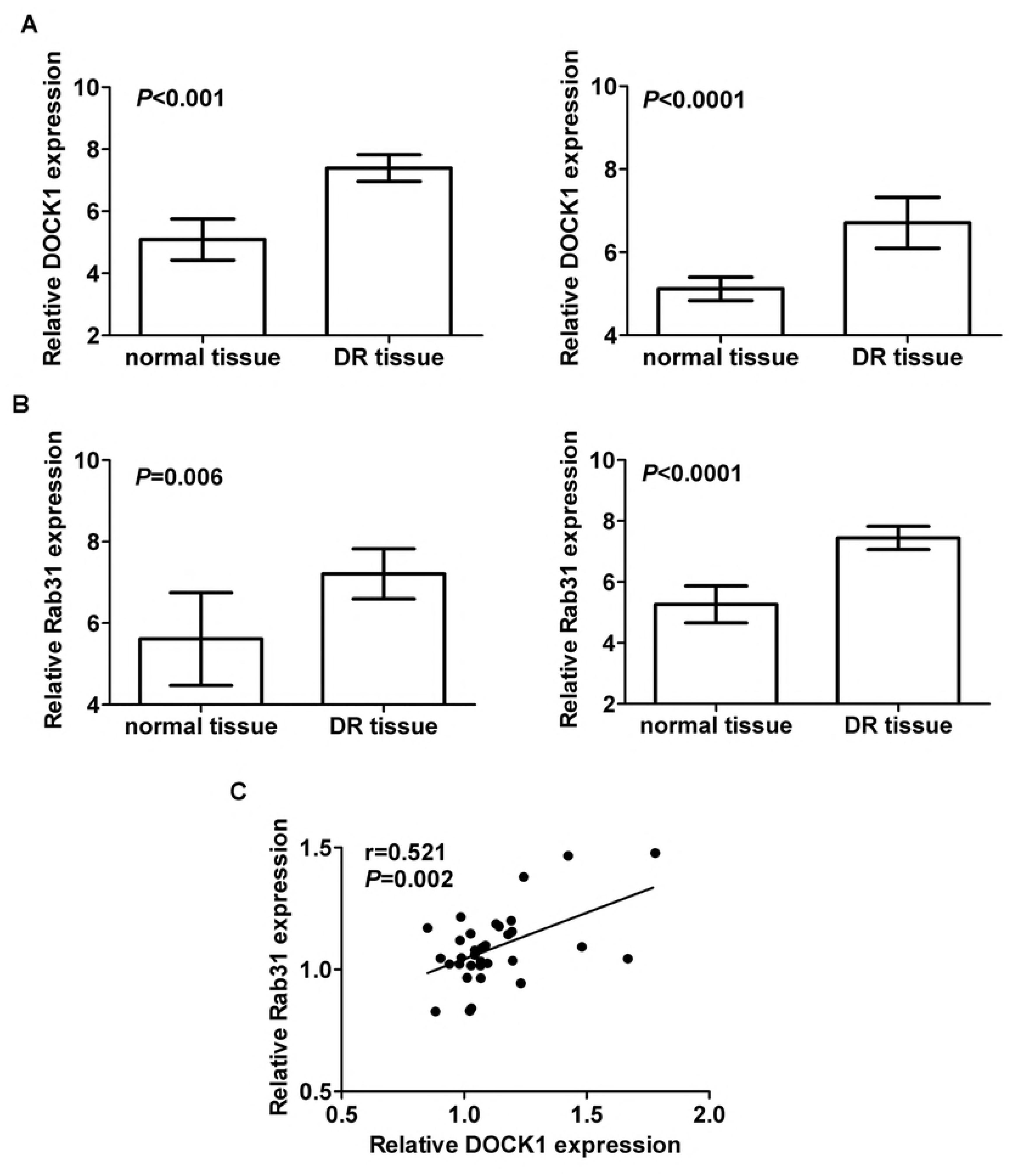
High expression and correlation of DOCK1 and Rab31 in DR. (A) Analysis of expression of DOCK1 and Rab31 (B) mRNA in fibrovascular membranes tissue of DR versus normal tissue and retinas from STZ induced diabetes mouse model. (C) Analysis of correlation of DOCK1 and Rab31 mRNA expression in four independent datasets, fibrovascular membranes tissue of DR from GEO (GSE53257 and GSE60436) and 16 retinas from STZ induced diabetes mouse samples from GEO (GSE24423 and GSE20886). All data are log transformed and median centered. Data were obtained from GEO public database. Data represent mean ± SEM. (\**P* < 0.05).

### 3.2 High glucose (HG) stimulation promotes proliferation of retinal epithelial cells and impaired tight junctions

Previous studies have shown that the indicator of HG-induced endothelial dysfunction is increased apoptosis [18]. To this end, we performed HG stimulation on retinal epithelial cells ARPE-19, and detected changes of TJ proteins expression and apoptosis. The results showed that the expression of TJ proteins (Occludin, Claudin-5 and ZO-1) in ARPE-19 cells was minimized (Figure 2A and B) at 30 nM HG-stimulated cells for 48h. Similarly, immunofluorescence experiments also confirmed this result (Figure 2C). At the same time, ARPE-19 cells maintained for 48h HG-stimulation showed a significant increase in cell apoptosis (Figure 2D) compared to cells maintained in normal glucose.

**FIG.2.**
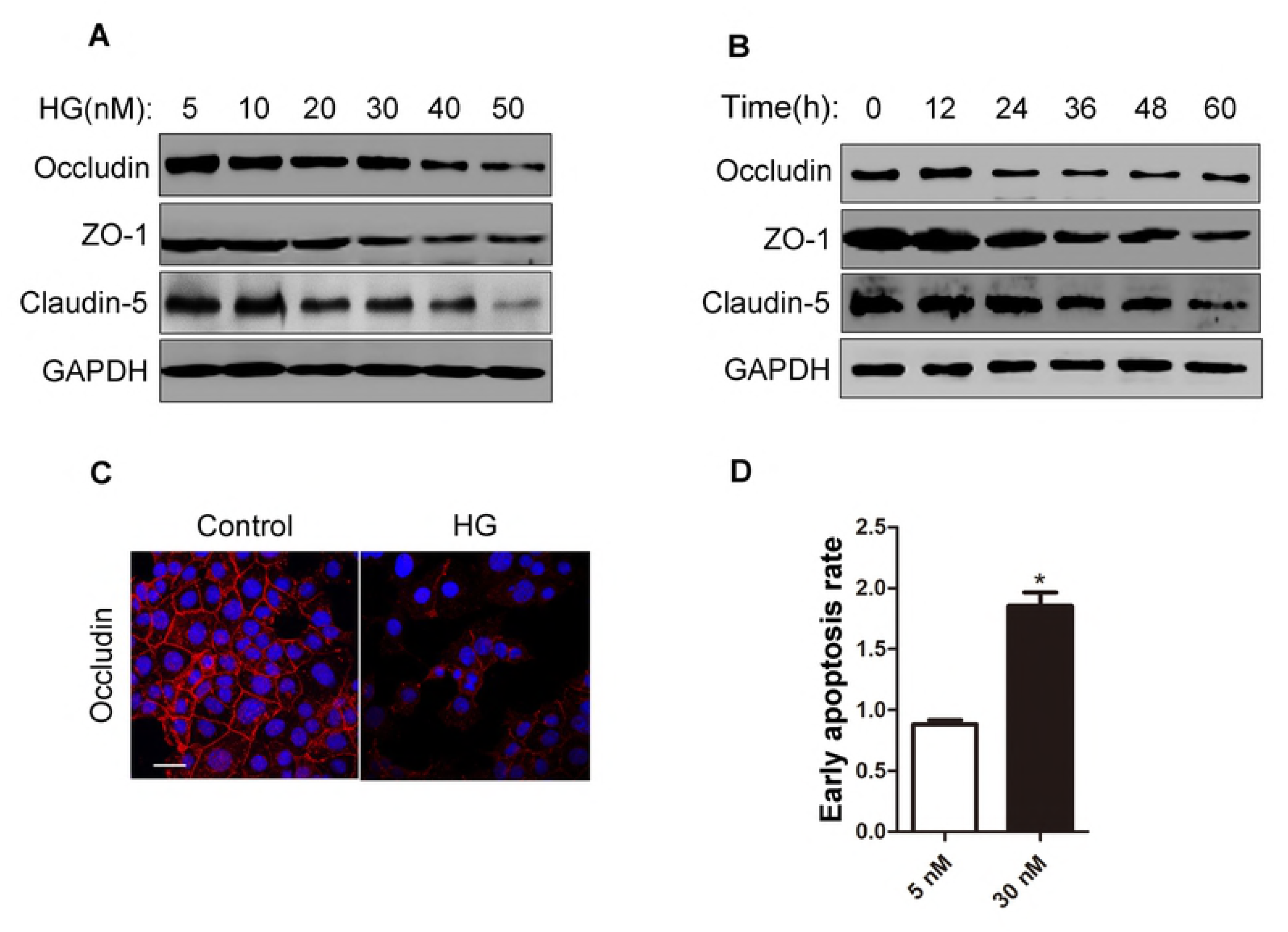
High glucose (HG) stimulation promotes proliferation of retinal epithelial cells and impaired tight junctions. (A) Western blotting detection for TJ proteins expression in ARPE-19 cells with different concentrations or (B) time with HG treatment. (C) After seeding ARPE-19 cells with or without HG treatment in 6-cm culture plates, the Occludin was immunofluorescent stain. (D) The cell apoptosis of ARPE-19 was detected using apoptosis detection assay after treatment with NG or HG treatment. Data represent mean ± SEM. (* *P* < 0.05).

### 3.3 HG stimulation promotes high expression of DOCK1 and Rab31 and activation of Rab31

RT-PCR showed that cells grown in HG medium showed a significant increase in mRNA levels of DOCK1 (Figure 3A) compared to ARPE-19 grown in normal medium. Immunoblot analysis also demonstrated that HG stimulation promoted high expression of DOCK1 at the protein level (Figure 3B). GAPDH expression used as an internal control confirmed that the amount of proteins added was equal in all groups. Furthermore, we also demonstrate that HG stimulation could increase the activity of Rab31. The HG treated group showed an increase in Rab31 activity by measuring the Rab31-GTP/total Rab31 ratio compared to the normal glucose (NG) treated group (Figure 3C). These results indicate that HG treatment not only increased the expression of DOCK1, but also increased the activity of Rab31.

**FIG.3.**
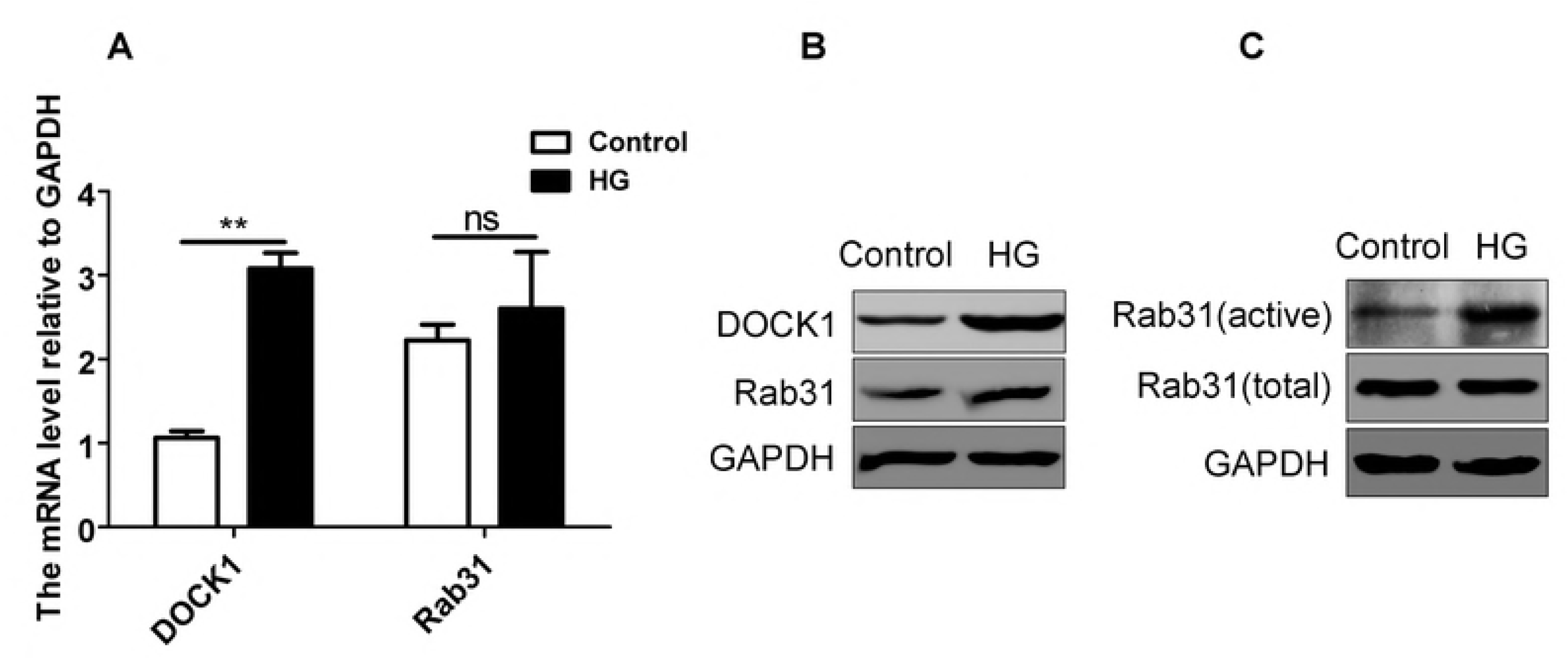
HG stimulation promotes high expression of DOCK1 and Rab31 and activation of Rab31. (A) The mRNA expression of DOCK1 and Rab31 in ARPE-19 cells with NG or HG treatment. (B) The expression of DOCK1 and Rab31 were detected by western blotting with NG or HG treatment.(C) The expression of active Rab31 and total Rab31 were detected by western blotting with NG or HG treatment after GST-pulldown assay. Data represent mean ± SEM. (** *P* < 0.01).

### 3.4 DOCK1 promotes the activation of Rac1

In view of above results, HG treatment can increase the expression of DOCK1, and DOCK1 acts as the GEF of Rac1. Therefore, we further verified whether the increased expression of DOCK1 further promoted the activation of Rac1. In ARPE-19 cells, compared with the normal HG-treated group, although the HG treatment could still slightly increase the activation of Rac1 after knockdown of DOCK1, the protein of Rac1-GTP was significantly decreased (Figure 4A). The slightly change in activity of Rac1 may be caused by other GEFs. At the same time, we have taken the way of overexpression to artificially increase the level of DOCK1 in the cells, and the activation of Rac1 is significantly increased compared with the control group (Figure 4B). In addition, in order to further study the effect of Rac1 on cell function, we examined changes in cell TJ and apoptosis. Compared with the control group, the use of siRNA to interfere with Rac1 expression did not prevent the change of Occludin upon HG stimulation (Figure 4C). However, HG-promoted apoptosis was significantly attenuated after knockdown of Rac1, suggesting that Rac1 mediates apoptosis in human retinal epithelial cells stimulated by HG (Figure 4D).

**FIG.4.**
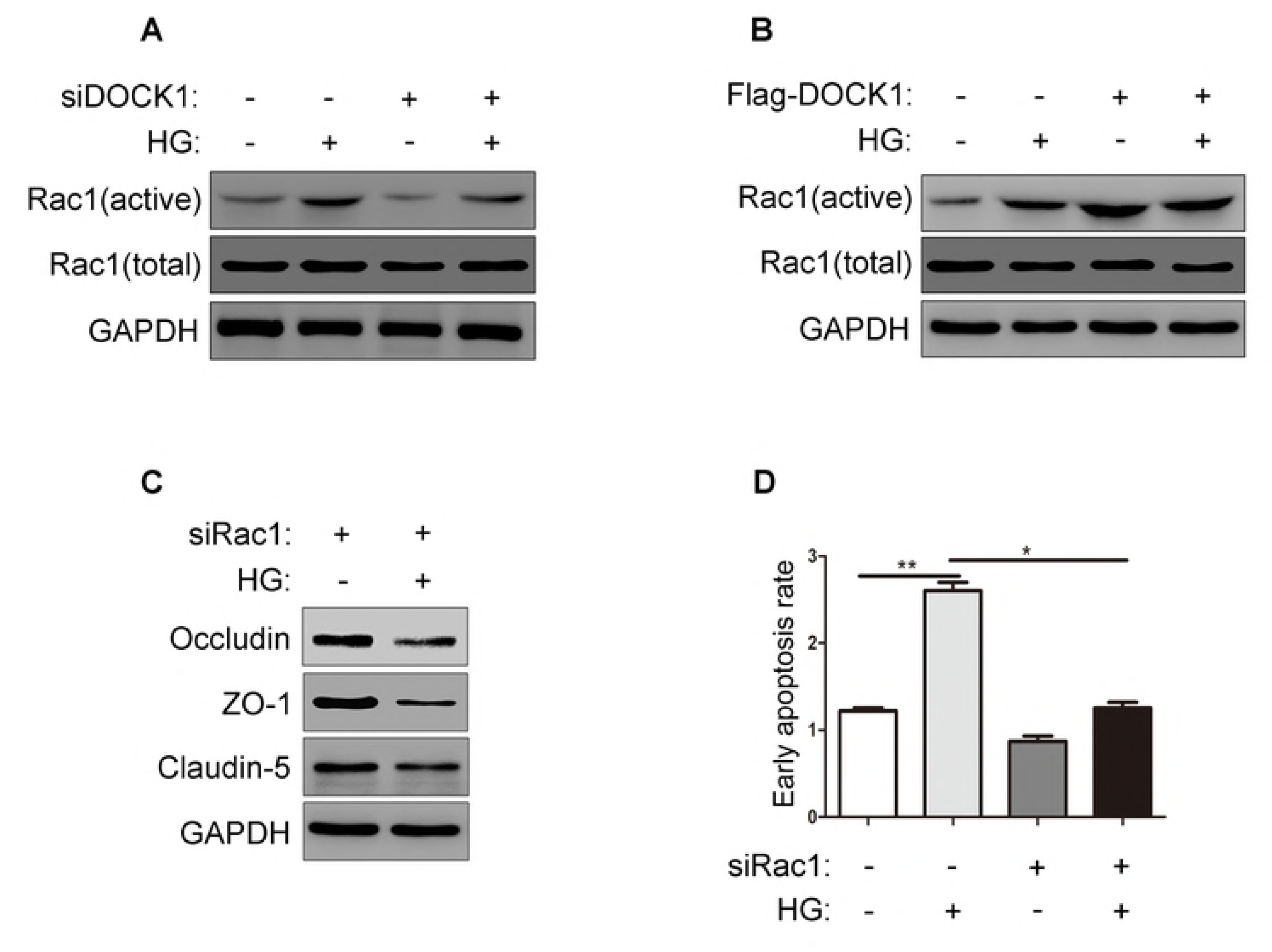
DOCK1 promotes the activation of Rac1. (A) ARPE-19 cells transfected with or without siDOCK1 for 24 h, and then the expression of active Rac1 and total Rac1 were detected by western blotting with NG or HG treatment after GST-pulldown assay. (B) ARPE-19 cells transfected with or without Flag-DOCK1 for 24 h, and then the expression of active Rac1 and total Rac1 were detected by western blotting with NG or HG treatment after GST-pulldown assay. (C) ARPE-19 cells transfected with or without siRac1 for 24 h, and then the expression of TJ proteins were detected by western blotting with NG or HG treatment. (D) The cell apoptosis of ARPE-19 transfected with or without siRac1 were detected using apoptosis detection assay after treatment with NG or HG treatment. Data represent mean ± SEM. (* *P* < 0.05, ** *P* < 0.01).

### 3.5 Activation of DOCK1-Rac1 signal promotes apoptosis

We further studied the effect of the DOCK1-Rac1 signal axis on apoptosis. It has shown that knockdown of DOCK1 could effectively inhibit apoptosis induced by HG (Figure 5A). In contrast, overexpression of DOCK1 further promotes HG-induced apoptosis (Figure 5B). Consistent with the above results, knockdown of Rac1 while overexpressing DOCK1 not only inhibited apoptosis upon HG stimulation, but also inhibited the increase of apoptosis induced by overexpression of DOCK1 (Figure 5C). However, overexpression of Rac1 while knockdown of DOCK1 still promotes increased apoptosis (Figure 5D). These results not only demonstrate the role of DOCK1-Rac1 signaling in apoptosis, but also demonstrate the signaling relationship between DOCK1 and Rac1.

**FIG.5.**
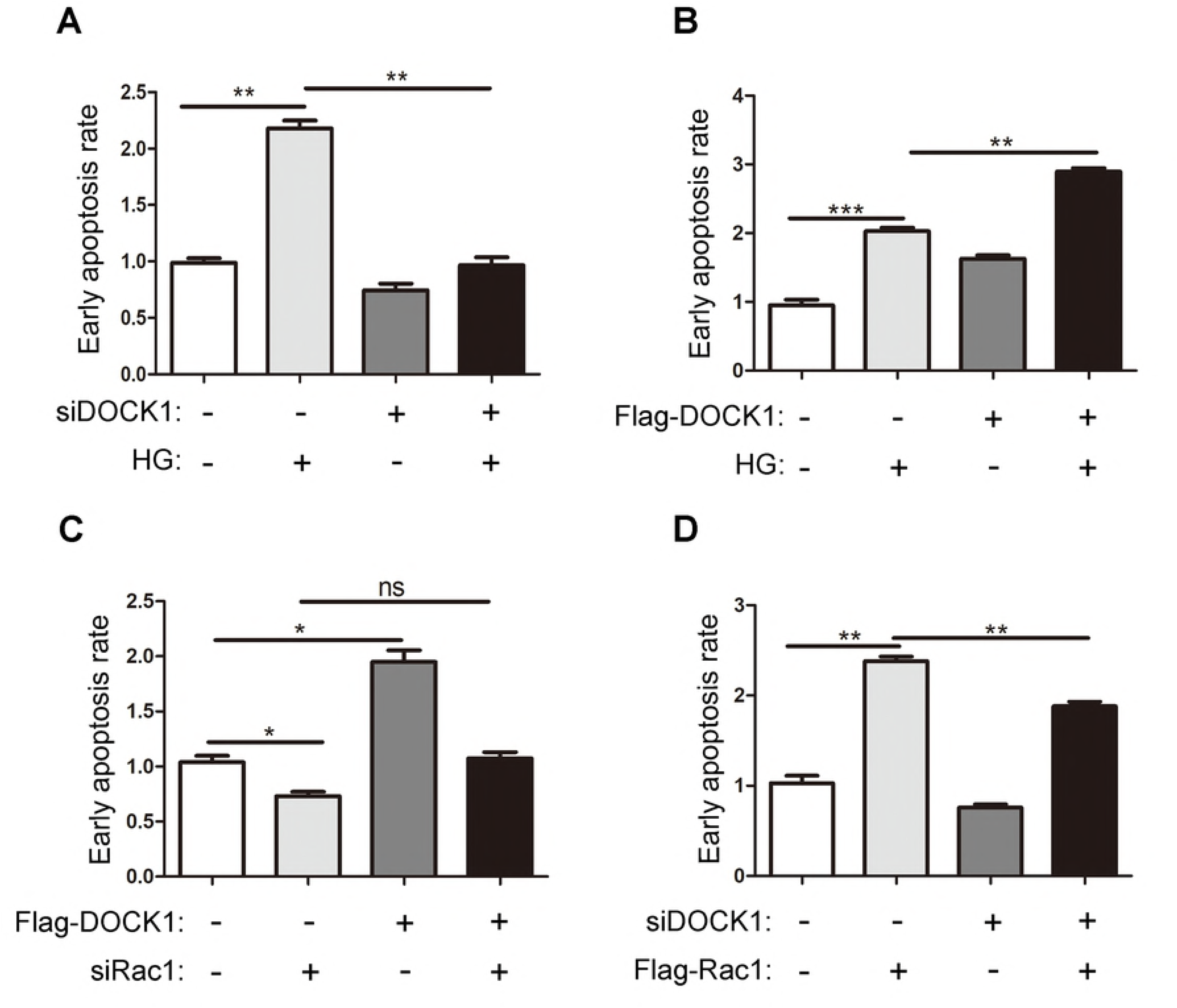
Activation of DOCK1-Rac1 signal promotes apoptosis. (A) The cell apoptosis of ARPE-19 transfected with or without siDOCK1 were detected using apoptosis detection assay after treatment with NG or HG treatment. (B) The cell apoptosis of ARPE-19 transfected with or without Flag-DOCK1 were detected using apoptosis detection assay after treatment with NG or HG treatment. (C) The cell apoptosis of ARPE-19 transfected with or without Flag-DOCK1 were detected using apoptosis detection assay after treatment with or without siRac1 transfection. (D) The cell apoptosis of ARPE-19 transfected with or without Flag Rac1 were detected using apoptosis detection assay after treatment with or without siDOCK1 transfection. (* *P* < 0.05, ** *P* < 0.01, \*\*\**P*<0.001)

### 3.6 High activation of Rab31 affects tight junctions of cells

As mentioned earlier, the classical TJ proteins consist of Claudin-5, Occludin and ZO-1. In our study, changes in the expression levels of these markers were also examined by western blotting. Compared with the control group, the expression level of Claudin-5, Occludin and ZO-1 was significantly decreased after 48 h of HG stimulation, while the knockdown of Rab31 could make this phenomenon disappear (Figure 6A). Similarly, the results of immunofluorescence using occludin as a marker also illustrate this (Figure 6B). To further explore the mechanism by which Rab31 is involved in and contribute to the reduction of TJ proteins, we performed colocalization studies using early endosomal markers (EEA1), late endosomal markers (M6PR), and lysosomal markers (LAMP1). After Rab31 activity was down-regulated by siRNA, the colocalization of Occludin with EEA1, M6PR and LAMP1 was significantly reduced (Figure 6C, D and E), indicating that Rab31 mediates endocytic transport of TJ proteins from plasma membrane to early endosome.

**FIG.6.**
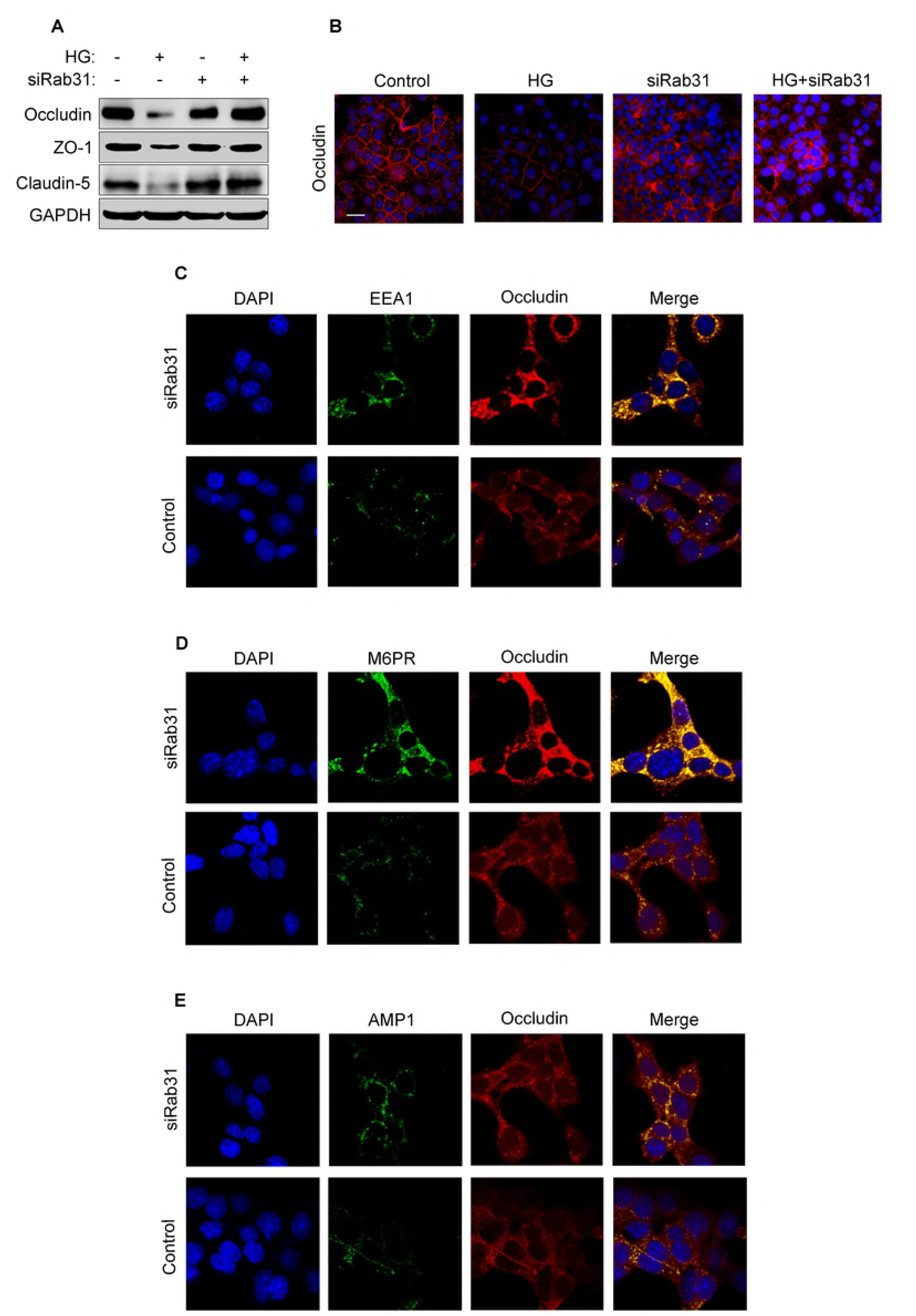
High activation of Rab31 affects tight junctions of cells. (A) ARPE-19 cells transfected with or without siRab31 for 24 h, and then the expression of TJ proteins were detected by western blotting with NG or HG treatment. (B) After seeding ARPE-19 cells with NG or HG treatment in 6-cm culture plates, the cells were transfected with or without siRab31. And then, Occludin was immunofluorescent stain in red and EEA1 (C), M6PR (D) and AMP1 (E) were immunofluorescent stain in green.

## 4. Discussion

DR is a major complication of diabetes and a major cause of vision loss caused by diabetes. In the early NPDR, it is characterized by retinal microvascular damage, which in turn leads to excessive blood vessel permeability and inflammation [19]. Although, the association between microvascular endothelial cell dysfunction (RMVED) caused by diabetes and diabetes has been confirmed in various studies [20]. Among them, some studies have mentioned that RhoA/ROCK1 signaling pathway and changes in TJ proteins play an important role in RMVED [21]. However, the specific molecular mechanism between the regulation of TJ proteins expression and even degradation has not been fully investigated in hyperglycemia-induced DR, especially the signal transduction pathways that may be involved.

Endothelial cell TJ proteins represent a structural barrier with selective paracellular permeability to solute and larger molecules, and its destruction enhances microvascular permeability [22]. There is increasing evidence that TJ is a dynamically regulated structure, and the dynamic changes in TJ also reflect the pathophysiological consequences of the disease. At the same time, such changes are also regulated by intracellular signaling pathways such as the Rab family of proteins. Occludin, Claudins and ZO-1 are major transmembrane proteins localized to TJ [23]. Studies have shown that Rab14 can degrade the TJ protein claudin-2 through interaction with PKC, thereby affecting the function of TJ [5]. Similarly, our study demonstrates that Rab31, which is highly expressed in the DR, especially high protein activity, promotes the degradation of TJ proteins in pathological conditions by mediating endocytic transport of TJ proteins from the plasma membrane to early endosomes. The high expression of Rab31 has been demonstrated in PDR vascular tissue, and the high activity of Rab31 was also found in the HG-stimulated DR model of this study. It has shown that high expression also means high activity, and the specific mechanism of the increased activity of Rab31 on the dynamic effects of TJ protein was confirmed.

Small GTPases play an important role in the development of many diseases [24]. In addition to playing a role in the regulation of a large number of signaling pathways in tumors, it has also been studied in DR. Studies have shown that the RhoA/ROCK1 signaling pathway can reduce the hyperosmoticity of endothelial cells caused by high glucose [21]. Studies have also shown that the Tima1-Rac1 signal axis is activated in the initial stages of diabetes, mediated activation of Nox2 and p38 MAP kinases to increase intracellular ROS, which not only leads to mitochondrial damage, but also accelerates capillary cell apoptosis, which in turn triggers early signaling events in the development of DR [12,13]. Tiam1 is a GEF of Rac1. Here, another GEF of Rac1, DOCK1, was studied. Database analysis demonstrated that DOCK1 is highly expressed in PDR vascular tissues. It has shown that the DOCK1-Rac1 signal axis promotes apoptosis in retinal epithelial cells in a HG-induced DR model. This phenomenon plays an extremely important role in retinopathy.

In conclusion, this study based on the HG-induced DR cell model reveals the role of two mutually synergistic signaling pathways through the important cytological phenomena of apoptosis and tight junctional damage. One is the degradation of TJ proteins caused by small GTPase Rab31-mediated vesicle trafficking, and the other is the apoptosis of retinal epithelial cells mediated by small GTPase Rac1 and its GEF DOCK1. Among them, the high expression of Rab31 and DOCK1 in PDR lesions showed a significant positive correlation. This also suggests a synergistic relationship between the two signaling pathways in this study. Clearer molecular mechanisms and interrelationships between signal pathways require more in-depth research through more living tissue and advanced experimental techniques.

